# Niche adaptation limits bacteriophage predation of *Vibrio cholerae* in a nutrient-poor aquatic environment

**DOI:** 10.1101/492439

**Authors:** Cecilia A. Silva-Valenzuela, Andrew Camilli

**Affiliations:** Department of Molecular Biology and Microbiology, Tufts University, Boston, Massachusetts, 02111, USA.

**Keywords:** *Vibrio cholerae*, bacteriophage, O1-antigen, fresh water, estuary

## Abstract

*Vibrio cholerae*, the causative agent of cholera, has reservoirs in fresh and brackish water where it interacts with virulent bacteriophages. Phages are the most abundant biological entity on earth and co-evolve with bacteria. It was reported that concentrations of phage and *V. cholerae* inversely correlate in aquatic reservoirs and in the human small intestine, and therefore that phages may quench cholera outbreaks. Although there is strong evidence for phage predation in cholera patients, evidence is lacking for phage predation of *V. cholerae* in aquatic environments. Here, we used three virulent phages, ICP1, ICP2, and ICP3, commonly shed by cholera patients in Bangladesh, as models to understand the predation dynamics in microcosms simulating aquatic environments. None of the phages were capable of predation in fresh water, and only ICP1 was able to prey on *V. cholerae* in estuarine water due to a requirement for salt. We conclude that ICP2 and ICP3 are better adapted for predation in a nutrient rich environment. Our results point to the evolution of niche-specific predation by *V. cholerae-*specific virulent phages, which complicates their use in predicting or monitoring cholera outbreaks as well as their potential use in reducing aquatic reservoirs of *V. cholerae* in endemic areas.

**Significance statement:** Virulent phages can reduce populations of bacteria and help shape bacterial evolution. Here, we used three virulent phages to understand their equilibrium with *V. cholerae* in nutrient-limiting aquatic microcosms. It has been proposed that phages quench cholera outbreaks, but no direct evidence of phage predation in aquatic environments had been established. Here we show that different phages possess varied abilities to infect in certain niches or stages of the host bacterial life cycle. Unveiling the phage/bacterial interactions in their natural setting is important to the understanding of cholera outbreaks and could be ultimately used to help develop a method for outbreak prediction and/or control.

## Introduction

*Vibrio cholerae* is a water-borne bacterium that causes cholera, an acute intestinal infection characterized by profuse secretory diarrhea that can quickly lead to severe dehydration and death if untreated. This pathogen is able to persist in aquatic environments in planktonic form or by association with phytoplankton and the chitinous carapaces of zooplankton (1) and is endemic in many countries where it forms reservoirs in fresh water and estuarine environments (2). Among the many interactions this pathogen could have in the environment, virulent phages may be of special relevance. Phages are viruses of bacteria that are present at ca. 10^30^ in the earth’s hydrosphere (3, 4). Phages exist in dynamic equilibrium with their bacterial hosts and help drive bacterial evolution and shape ecosystems (3, 5).

In the 1920’s Felix D’Herelle studied the relationship between *V. cholerae* and virulent phages, finding that phages were widespread in the environment a few days after the onset of an outbreak (6). Recent studies have proposed that cholera outbreaks are driven in part by an increase in the hyperinfectious *V. cholerae* population (7, 8), and quenched by an increase of virulent phages in infected humans (9, 10) and in environmental reservoirs (11, 12). However, no direct evidence of phage predation of *V. cholerae* in aquatic settings has been reported. It is known that phage can limit *V. cholerae* replication during intestinal infection (13) and are shed into the environment due to fecal contamination of waters in endemic areas. Hence, the appearance of environmental phage may simply be a consequence of phage replication in infected humans (9, 10).

Virulent phages are frequently found in cholera patient stools (14, 15). A Study of a 10-year collection of such stool samples in Bangladesh revealed the presence of just three *V. cholerae-*specific virulent phages, ICP1, ICP2, and ICP3 (15). Of these, ICP1 was the most frequently observed. Because of the dominance of these three phages in this area, the fact that they are often shed at high titers in secretory diarrhea, and the general lack of waste-water treatment facilities, it is likely that these phages frequently contact *V. cholerae* in the surrounding aquatic environments.

We used ICP1, ICP2, and ICP3 to study their predation abilities on *V. cholerae* in microcosms simulating fresh and estuarine aquatic environments. Our results show that none of the phages are able to prey on *V. cholerae* in fresh water, and only ICP1 is able to prey in estuarine water. ICP1 can replicate in *V. cholerae* in fresh water that is supplemented with NaCl, hence, salinity could be a critical variable in relation to environmental phage predation.

## Results

### ICP1 preys on planktonic *V. cholerae* in estuary conditions

Cholera epidemics occur in Bangladesh during March-May and after monsoon season in September-December. The temperature of bodies of water in Bangladesh approaches 30°C (16). Therefore, we first tested the ability of these phages to kill *V. cholerae* in a rich (Luria-Bertani [LB]) broth medium at 30°C. When added to a multiplicity of infection (MOI) of 0.01, all three phages were able to reduce the load of *V. cholerae* over the first few hours. ICP1 and ICP3 reduced bacterial numbers 100-fold within 1h, but after 4h, numbers increased due to the outgrowth of phage-resistant (escape mutant) bacteria (SI appendix, **Figure S1A**). ICP2 infected cultures kept growing for 1h, but by 3h bacterial numbers were reduced 10,000-fold. Therefore, each of the ICP phages can infect and kill *V. cholerae* at 30°C. This temperature was used throughout the rest of this study.

It has been shown that upon dissemination of *V. cholerae* from cholera patient stools to fresh water there is a major drop in osmolarity, inorganic nutrients and carbon sources (10). An obstacle to the study of phage predation in an aquatic model is the loss of bacterial viability in this nutrient-poor environment (10, 17). Since *V. cholerae* shed from the host resembles the physiology of stationary growth phase bacteria (18) and because stationary phase cells exhibit better survival to osmotic shock (17), we used stationary phase cultures for adaptation to the aquatic environment prior to phage infection.

Phages use the bacterial cell machinery in order to replicate and produce progeny (19). Since shed *V. cholerae* resemble stationary phase bacteria, we tested if any of the phages were able to kill *V. cholerae* in this physiological state. 10^6^ colony-forming units (CFU) of *V. cholerae* from an overnight, stationary phase culture were inoculated into M9 minimal media in the absence of a carbon source to avoid bacterial growth and phages were added to an MOI of 3. ICP1 was able to reduce the *V. cholerae* load within 1h by 10-fold. ICP1 killing was sustained for 3h, but after 4h, bacterial numbers increased due to the generation of ICP1 escape mutants (SI appendix, **Figure S1B**). ICP2 was able to kill *V. cholerae* by 3-fold within 1h but then bacterial growth quickly recovered. ICP3 was not able to kill *V. cholerae* under these conditions.

To study phage predation on planktonic *V. cholerae* in water in the absence of added carbon source, 10^7^ CFU of stationary phase *V. cholerae* were pre-adapted overnight at 30°C with aeration in either 0.7% Instant Ocean, which we chose to mimic estuary, or fresh water obtained from a natural pond. For comparison, open ocean is equivalent to 3.5% Instant Ocean. Cultures were then infected with each phage at an MOI of 0.01 and bacterial viability was measured over time. Only ICP1 was able to replicate and kill *V. cholerae* in estuary conditions (**Figure 1A, Figure S2A and B**). The *V. cholerae* population was reduced 10-fold by 4h and 10,000-fold by 7h. After 24h, viable bacterial counts rebounded by means of the growth of escape mutants. All phages were able to adsorb to *V. cholerae* in this environment (**Figure 1B**) and were stable at 30°C with aeration in the absence of a bacterial host (SI appendix, **Figure S3A**), thus ruling out these trivial explanations for the lack of predation by ICP2 and ICP3.

**Figure 1.**
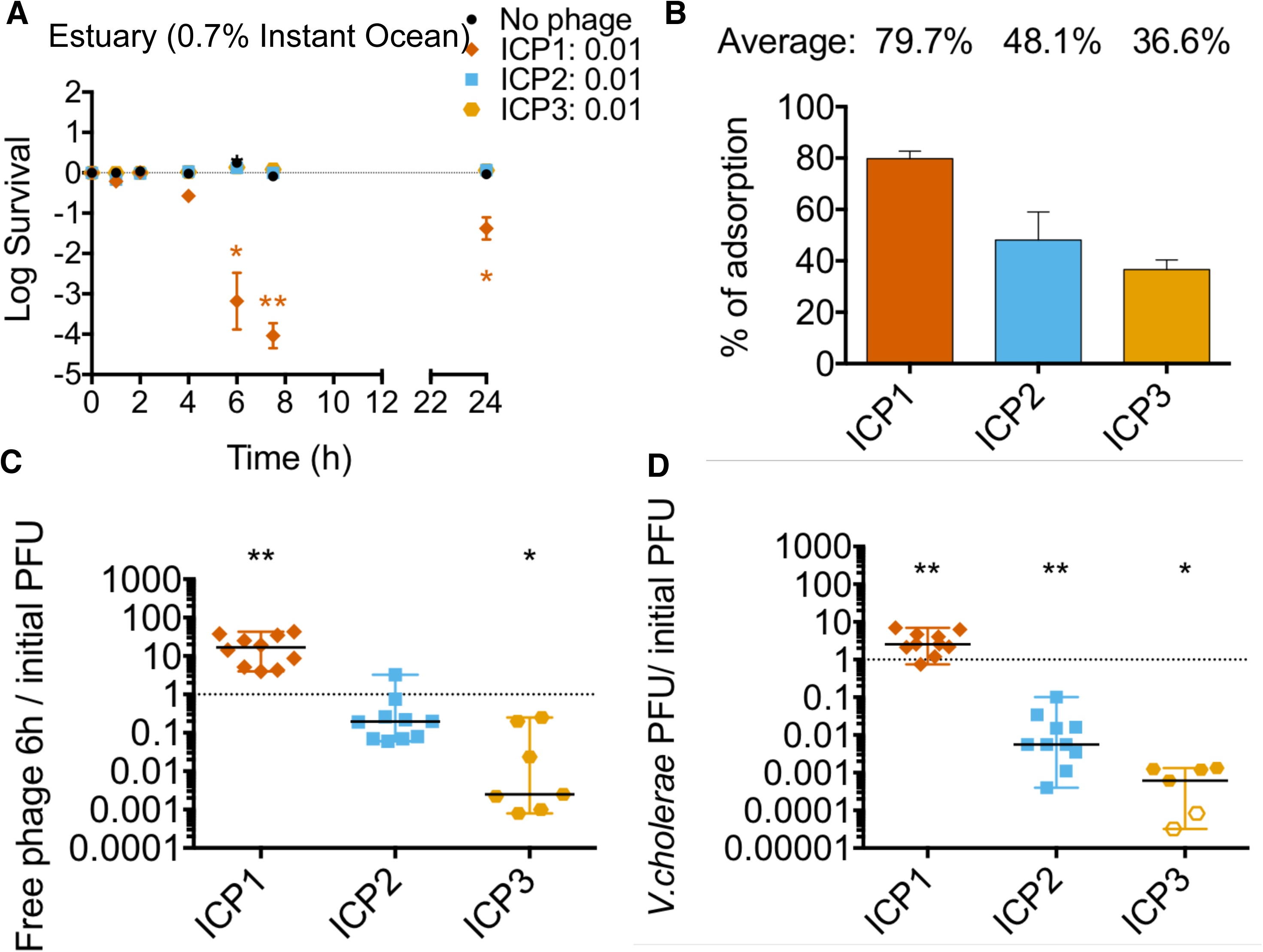
ICP1 preys on *V. cholerae* in estuary water. 10^7^ CFU of *V. cholerae* from an overnight culture were pre-adapted in 0.7% Instant Ocean overnight in the absence of added carbon source. ICP1, ICP2 or ICP3 were added to an MOI of 0.01 to assess bacterial viability and phage replication over time. Panels a and b show average with estandar error (a) Bacterial viability at 1, 2, 4, 7 and 24h. (b) Phage adsorption after 15 min of static incubation at 24°C. Panels c and d show the medians and range of at least eight biological replicates. Empty symbols represent data under the limit of detection. (c) Ratio of phage in culture supernatant at 6h relative to number initially added. (d) Ratio of phage associated with *V. cholerae* at 6h relative to number initially added. (*P<0.05, **P<0.005).

The number of free and cell-associated phage after 6h of infection was determined by infectious-centre (plaque) assay. To measure cell-associated phage, the cells were washed prior to performing plaque assays. Removal of greater than 99% of unbound phage was confirmed by quantifying free phage before and after washing (**Figure S2C**). Consistent with its ability to kill *V. cholerae* in estuarine conditions, the titer of ICP1 in the culture supernatant increased by 15-fold (**Figure 1C**). In contrast, the titers of ICP2 and ICP3 in the culture supernatant decreased 1- and 3-orders of magnitude, respectively. In terms of cell-associated phage at 6h, the titer of ICP1 increased 3-fold relative to the amount of phage added initially (**Figure 1D**). In contrast, only 1% of ICP2 and 0.1% of ICP3 were cell-associated (**Figure 1D**). Thus, the majority of bound ICP2 and overwhelming majority of ICP3 were unable to replicate in estuary-like conditions and lost their ability to subsequently form plaques.

Altogether, these results suggest that in estuarine conditions where *V. cholerae* is in a non-growing planktonic state: (i) ICP1 is efficient at adsorption, predation and replication, (ii) ICP2 is able to adsorb but only a small percentage of these can replicate with most of the remaining losing plaque-forming ability, and (iii) ICP3 is able to adsorb but cannot replicate nor maintain infectivity.

### No ICP phage can prey on *V. cholerae* in fresh water

It has been suggested that fresh water is the primary aquatic reservoir for water-borne transmission of *V. cholerae* during epidemics (20). Therefore, understanding the phage predation dynamics in this environment is crucial. For this, we performed the same set of predation experiments as above, but using fresh water obtained from a natural pond. Our results show that none of the ICP phages were able to kill *V. cholerae* under these conditions (**Figure 2A**). Since lack of predation did not seem to be due to either lack of adsorption (**Figure 2B**) or phage stability (SI appendix, **Figure S3B**), this strongly suggests that phage predation/replication does not occur in fresh water environments. Notably, ICP1 and ICP2 had a higher adsorption rate in estuary conditions (compare **Figures 1B** to **2B**). In fresh water, ICP1 and ICP2 were found mostly as free phage in the supernatant after 6h and only 1%-2% were either associated with, or within infected bacterial cells (**Figure 2C** and **2D**). As seen previously, less than 1% of the total ICP3 added was found in the supernatant after 6h, but less than 0.001% was able to replicate after interacting with cells.

**Figure 2.**
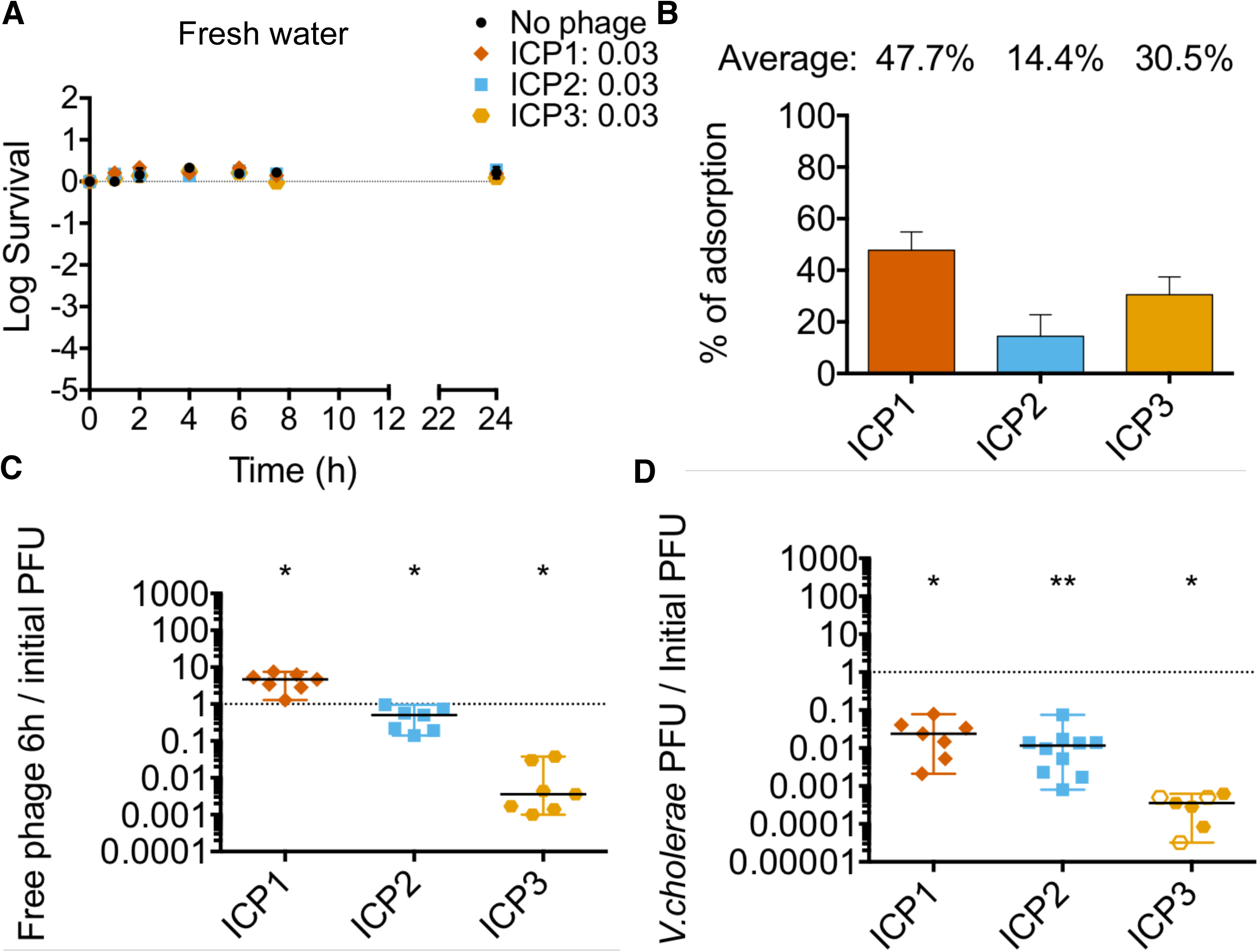
None of the phages preys on *V. cholerae* in fresh water. 10^7^ CFU of *V. cholerae* from an overnight culture were pre-adapted in autoclaved, filter-sterilized fresh water overnight in the absence of added carbon source. ICP1, ICP2 or ICP3 were added to an MOI of 0.01 to assess bacterial viability and phage replication over time. Panels a and b show average with estandar error (a) Bacterial viability at 1, 2, 4, 7 and 24h. (b) Phage adsorption after 15 min of static incubation at 24°C. Panels c and d, show the median with range of at least six biological replicates. Empty symbols represent data under the limit of detection. (c) Ratio of phage in culture supernatant at 6h relative to number initially added. (d) Ratio of phage associated with *V. cholerae* at 6h relative to number initially added. (*P<0.05, **P<0.005).

Together the results suggest that: (i) ICP1 can kill *V. cholerae* and replicate in a higher salinity condition than the one found in fresh water, (ii) only a small number of ICP2 phages replicate in *V. cholerae* in an estuary environment, and (iii) infection by ICP3 is unsuccessful in both aquatic environments.

### ICP1 predation in estuary water selects for escape mutants

We observed that between 7 and 24h post infection in estuary conditions, bacterial viability increased in the presence of ICP1 (**Figure 1A**) and the likely explanation for this is the generation of escape mutants via phase variation which alters expression of the lipopolysaccharide O1-antigen phage receptor (21). We tested this by plating surviving *V. cholerae* at 7 and 24h post infection, testing the isolates for phage sensitivity by cross-streaking, and then performing whole genome sequencing to identify mutations. The number of escape mutants increased over time after 7h of infection (SI appendix, **Figure S4A**). A total of 8 escape mutants after 24h of infection were sequenced, and mutations were identified in O1-antigen biosynthetic, phase variable genes *wbeL* and *manA* (21), as well as in other LPS biosynthetic genes (SI appendix, **Figures S4B, S4C and supplementary table 1**). At this same time, a small percentage of survivors (4%) were still sensitive to infection by ICP1 (SI appendix, **Figure S4A,** 24h). No ICP2 or ICP3 escape mutants were detected in the estuary cultures, consistent with the lack of phage replication (**Figure 2A**).

### Salinity is a crucial factor for ICP1 predation

Since ICP1 was able to kill *V. cholerae* in estuary-like conditions and not in fresh water, we hypothesized that either the physiology of *V. cholerae* varies in the two environments or that a co-factor present in estuary but not in fresh water might be needed by ICP1 for predation. One major difference between estuary and fresh water is salinity. The concentration of Na^+^ in fresh water is about 0.3 mM (10) compared to 92.4 mM in 0.7% instant ocean (22). To test if ICP1 replication requires high salinity, NaCl was added to fresh water to the levels present either in our estuarine condition (0.09 M) or LB broth (0.17 M). Cultures of *V. cholerae* pre-adapted in the two NaCl concentrations tested were infected using a MOI of 0.01. ICP1 was able to infect and kill *V. cholerae* in fresh water supplemented with 0.09 or 0.17 M NaCl by 7h, albeit 100-fold less than in estuary conditions (**Figure 3A**). Consistent with these levels of killing, we observed moderate levels of phage replication compared to the results from estuary water (**Figure 3B**). These results suggest that NaCl alone has an impact in predation by ICP1, however, this was not sufficient to obtain the same level of predation seen in estuary conditions. Fresh water often contains growth limiting amounts of nutrients such as phosphate and fixed nitrogen (10) and this may prevent *V. cholerae* from fully supporting ICP1 infection.

**Figure 3.**
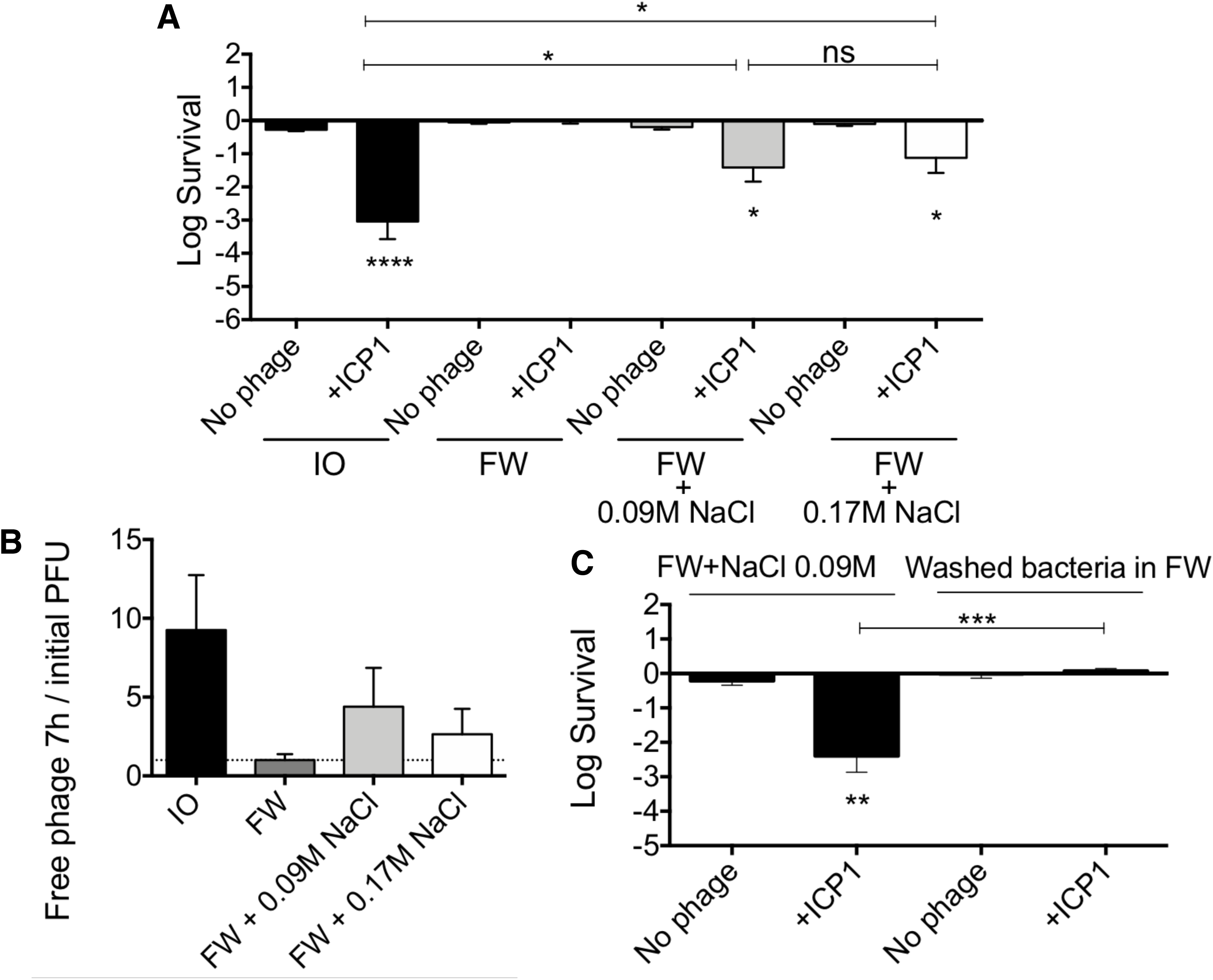
A salinity threshold is needed for ICP1 predation. 10^7^ CFU of *V. cholerae* from an overnight culture were pre-adapted overnight in 0.7% Instant Ocean, fresh water or fresh water supplemented with NaCl (0.09M or 0.17M) in the absence of added carbon source. ICP1 was added to an MOI of 0.01 to assess bacterial viability and phage replication over time. Shown is the average with standard error of at least six biological replicates. (a) Bacterial viability at 7h. (b) Ratio of phage in culture supernatant at 7h relative to number initially added. (c) Bacterial viability at 7h. (*P<0.05, **P<0.005, ****P<0.0001).

Because *V. cholerae* is a halophile, growth in high salinity may be required for the cells to become susceptible to ICP1. Alternatively, ICP1 attachment or infection may require the presence of salt. To differentiate between these, we pre-adapted *V. cholerae* in fresh water supplemented with 0.09 M NaCl then washed and resuspended the cells in fresh water alone. If NaCl is necessary for phage attachment or infection, changing the media to fresh water alone should impair predation. Cultures were washed and infected with ICP1 at an MOI of 0.01 and viable CFU were assessed at 7h. ICP1 exhibited impaired predation (**Figure 3C**) similar to the levels seen in fresh water alone (compare **Figure 2B** and **3A**). These results indicate that, at the time of infection, ICP1 predation in an aquatic condition requires a minimum salinity threshold well above the salinity of fresh water.

### Chitin availability aids phage predation

It has been proposed that *V. cholerae* can grow in the environment using chitin as a nitrogen and carbon source (23). We first tested whether *V. cholerae* could utilize chitin for growth in our aquatic conditions by adding 10^4^ CFU from a stationary phase culture into a 1% chitin suspension in either 0.7% Instant Ocean or fresh water. *V. cholerae* was able to grow in both, however, growth in the estuarine condition is faster and reaches a higher yield when compared to fresh water (SI appendix, **Figure S5**).

To evaluate if chitin utilization aids phage predation, we tested *V. cholerae* both in exponential and stationary phases of growth. For exponential growth in the estuary condition plus chitin, pre-adaptation was done for 8h at 30°C in static conditions (SI appendix, **Figure S5B**). Each phage was then added to an MOI of 0.01 and incubated for 16h, after which bacterial viability and phage replication was assessed. ICP1 reduced the *V. cholerae* population only by 10-fold (**Figure 4A**) due to the generation of escape mutants during the incubation period. On the other hand, neither ICP2 nor ICP3 caused a reduction in bacterial numbers. Consistent with predation by ICP1, the phage titer increased 10,000-fold, while ICP2 and ICP3 titers showed no significant increase (**Figure 4B**).

**Figure 4.**
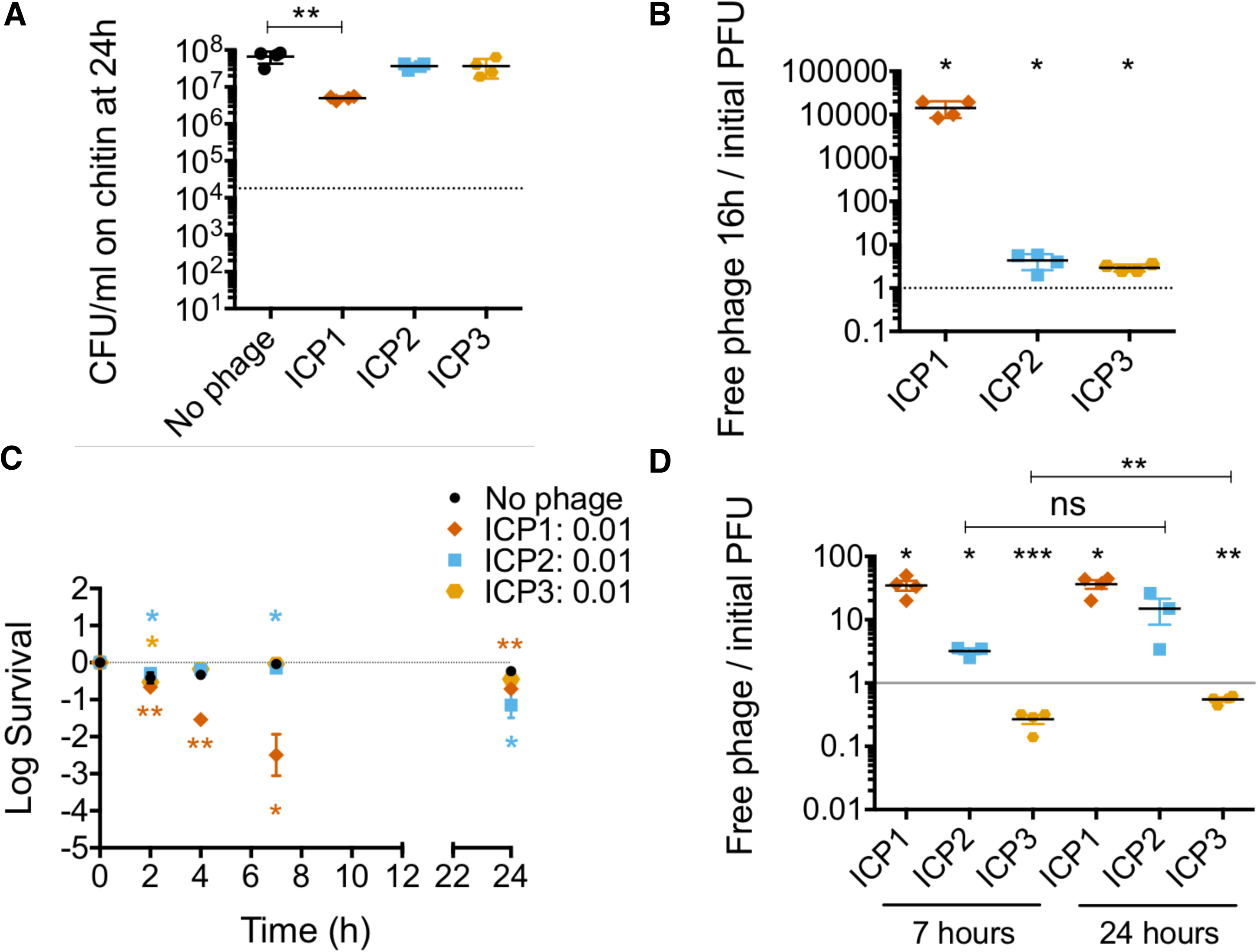
*Vibrio cholerae* is more susceptible to ICP1 and ICP2 in the presence of chitin. 10^4^ CFU of *V. cholerae* from an overnight culture were pre-adapted in a 1% chitin suspension in 0.7% Instant Ocean until exponential growth. ICP1, ICP2 or ICP3 were added to an MOI of 0.01 to assess bacterial viability and phage replication over time. Shown is the average with standard error of at least three biological replicates. (a) Bacterial viability at 16h. (b) Phage replication at 16h relative to number initially added. (c) 10^7^ CFU of *V. cholerae* from an overnight culture were pre-adapted overnight for phage predation. ICP1, ICP2 or ICP3 were added to an MOI of 0.01. Bacterial viability at 1, 2, 4, 7 and 24h in estuary conditions is shown. (d) Ratio of phage in culture supernatant at 7h and 24h relative to number initially added. (*P<0.05, **P<0.005, ****P<0.0001).

To test stationary phase cells, 10^7^ CFU from an overnight culture were inoculated in a 1% chitin suspension in either 0.7% instant ocean or fresh water and cultures were incubated overnight at 30°C in static conditions. Each phage was then added to a MOI of 0.01 and bacterial viability and phage replication were measured. In fresh water none of the phages were able to kill *V. cholerae* (SI appendix, **Figure S6A**) or replicate (SI appendix, **Figure S6B**). In the estuary environment, we observed results similar to the estuary planktonic conditions. Only ICP1 had the ability to kill *V. cholerae* and to replicate (**Figure 4C** and **4D**). Consistent with this killing, escape mutants arose; after 7h of infection, 90% of the survivors were resistant to infection by ICP1 and this number grew to 98% by 24h (SI appendix, **Figure S4D**). ICP2 exhibited a slightly increased ability to kill *V. cholerae* and to replicate compared to estuary alone; Viability of *V. cholerae* was decreased by 10-fold at 24 h and phage titer increased 10-fold. Consistent with this moderate level of killing, at 7h of infection 1 out of 24 isolates was resistant to ICP2 (SI appendix, **Figure S4E**). In contrast, and similar to that observed in estuary alone, ICP3 was not able to reduce *V. cholerae* viability nor replicate (**Figure 4C** and **4D**). Since *V. cholerae* was able to grow on chitin in both aquatic conditions (SI appendix, **Figure S5A**), the lack of phage predation in fresh water is not due to a lack of bacterial growth but instead appears to be impaired by the physicochemical properties of the water and/or the host’s response to growth in those conditions.

### ICP2 and ICP3 bound to *V. cholerae* do not replicate in the infant mouse small intestine

ICP1 replication in high salinity aquatic environments seems plausible according to our results, and could explain the high prevalence of ICP1 in Bangladesh. However, it is unclear how ICP2 and ICP3 are maintained. ICP2 and ICP3 have only been observed to naturally replicate in *V. cholerae* that infects the intestine and the data provided here raise doubt as to their ability to replicate on *V. cholerae* in nutrient poor environmental waters. Since both phages and bacteria are shed following human infection, it is possible that ingestion of contaminated water could perpetuate phage replication. One question that arises is whether phage and bacteria are ingested as separate entities or whether they are co-associated. The former has already been demonstrated in two different animal models of intestinal infection, wherein phage added prior to challenge with *V. cholerae* or concurrently substantially decrease intestinal colonization (9, 12).

To test the latter hypothesis, we first examined the ability of cell-associated phages in fresh water to resume replication upon a shift to a nutrient rich condition. We infected 10^7^ CFU in fresh water with each phage for 45 min at an MOI of 1 in static conditions at 24°C. Cultures were washed to remove free phage, and the cells carrying bound phage were added to 2 ml of LB broth and grown (outgrowth) at 37°C with aeration. After the wash step, 20% of ICP2 added and 0.01% of ICP3 added were cell-associated and yielded live virions, similar to the planktonic predation results shown previously (**Figure 1D**). Subsequently, both ICP2 and ICP3 reduced *V. cholerae* numbers after 3h of outgrowth (**Figure 5A**). This indicates that, when the small number of bacteria with live bound phage resumed growth, both ICP2 and ICP3 were able to resume replication and kill the bacteria. By 6h, bacterial numbers increased by means of the growth of escape mutants. Consistent with phage predation, ICP2 titer increased by 100-fold at 3h and by 1000-fold at 6h (**Figure 5B**); ICP3 titer increased by 5000-fold by 3h (**Figure 5B**). Also consistent with phage predation, at 6h of outgrowth, 100% (24/24) of the *V. cholerae* isolates were ICP2 escape mutants (**Figure 5C)**. For the ICP3 infected cultures at 6h, 75% (22/29) of *V. cholerae* isolates recovered were escape mutants (**Figure 5C**).

**Figure 5.**
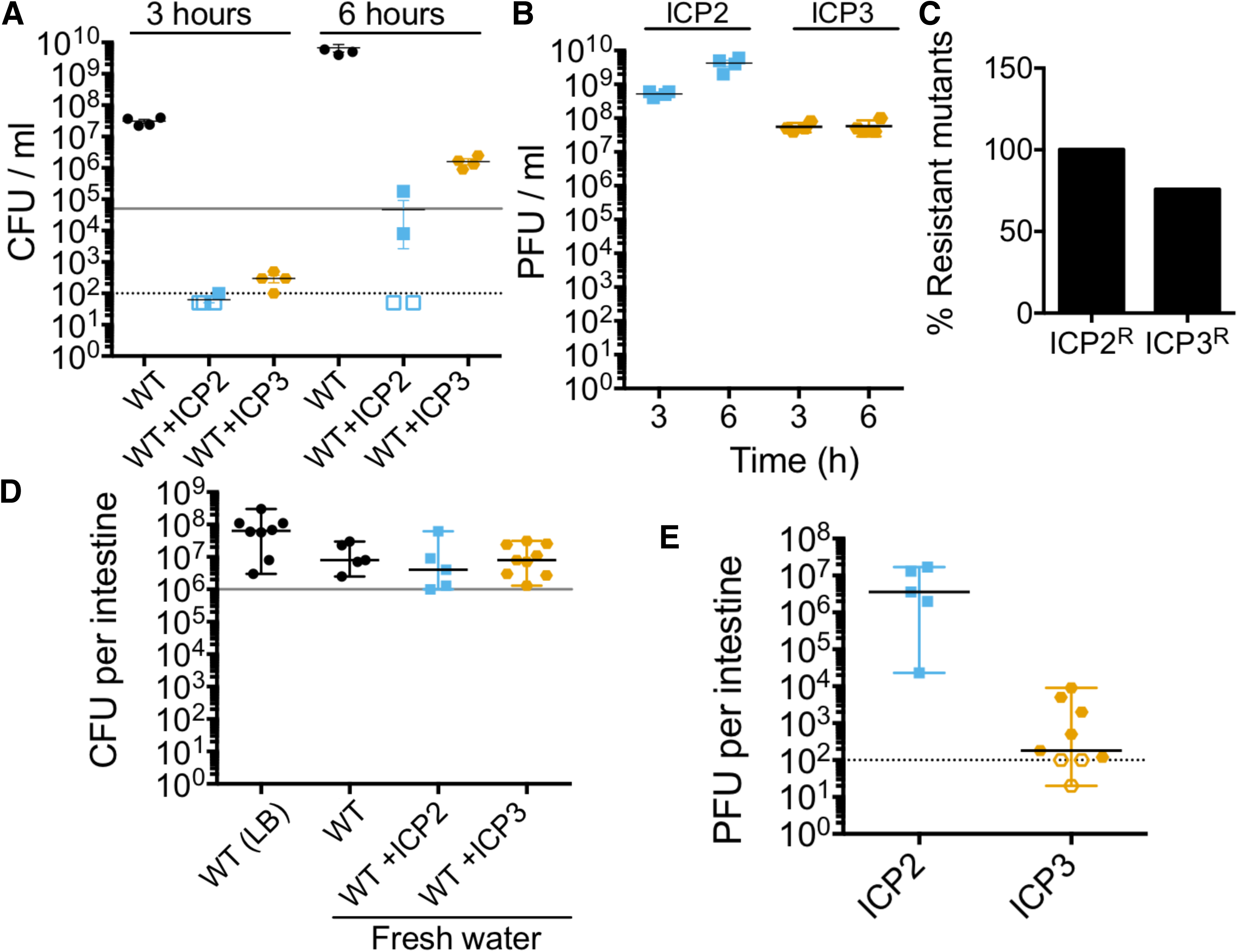
ICP2 and ICP3 replicate in a nutrient-rich environment. 10^7^ CFU of *V. cholerae* from an overnight culture were pre-adapted in fresh water overnight in the absence of added carbon source. ICP2 or ICP3 was added to an MOI of 1 and adsorption was conducted for 45 min. Cultures were washed and inoculated into 2 ml of LB broth or into CD-1 mice for assessment of bacterial colonization and phage replication. Empty symbols represent data under the limit of detection. (a) Bacterial replication in LB broth at 37°C with aeration at 3 or 6h of outgrowth of four independent replicates. Grey line indicates bacterial concentration at time 0. (b) Phage titer from LB cultures at 3 or 6h of outgrowth. (c) Average of percentage of resistant mutants for ICP2 and ICP3 at 6h of outgrowth. (d) Bacterial colonization of the small intestine in CD-1 infant mice. Graph represents the median with range of at least five biological replicates. Grey line indicates bacterial concentration in the inoculum. (e) Phage titer from colonized small intestine. Graph represents the median with range of at least five biological replicates.

We next examined the ability of bound phages to reduce *V. cholerae* colonization of the small intestine of 5-day-old mice. 10^6^ CFU of stationary phase *V. cholerae* that were pre-adapted overnight in fresh water were infected with ICP2 or ICP3 for 45 min in fresh water at an MOI of 1. Cultures were washed and resuspended in fresh water for orogastric inoculation into mice. As controls, mice were also inoculated in the absence of phage with 10^6^ CFU of *V. cholerae* grown overnight in LB broth or pre-adapted overnight in fresh water. After 24h of infection, mice were euthanized, the small intestines were dissected, homogenized, and the titers of *V. cholerae* and phage were determined. Pre-adaptation overnight in fresh water did not reduce the colonization ability of *V. cholerae*, since the load of bacteria per small intestine was not different from that of *V. cholerae* grown in LB (**Figure 5D**). Our results show that cell-associated ICP2 and ICP3 do not limit *V. cholerae* infection, since the *V. cholerae* loads present in those animals were not different from either of the control animals (**Figure 5D**).

ICP3 was undetectable in a third of the animals, and observed at low titer in the remaining ones (**Figure 5E**). The inability of ICP3 to limit *V. cholerae* colonization of mice under these conditions can be explained by its inability to replicate after attachment to cells in fresh water. In contrast, ICP2 appeared to be able to replicate at least to some extent, since all of the small intestines had a phage titer slighly higher than the number inoculated (**Figure 5D**). However, this was not sufficient to reduce *V. cholerae* colonization of these animals. Consistent with this, no *V. cholerae* escape mutants to ICP2 were found among 40 isolates screened (8 from each mouse output). Thus, cell-associated ICP2 and particularly ICP3 fare poorly when inoculated into the infant mouse intestinal tract and, as a result, are not able to reduce the load of *V. cholerae* to any measurable extent.

## Discussion

A better understanding of the ecology of *V. cholerae* and its phages in aquatic reservoirs could lead to the development of better tools for outbreak prediction and potential treatments to water supplies to reduce human infection. We used fresh and estuary water microcosms to study the interaction of three virulent phages with *V. cholerae*. These experiments are reductionist, since in its reservoir *V. cholerae* has complex interactions with other organisms and it also experiences changes in the physicochemical properties of the water throughout the year (24). Nevertheless, our results have relevance for understanding the impact of virulent phages on *V. cholerae* during cholera outbreaks, during which *V. cholerae* and phages are shed in secretory diarrhea into bodies of water and are subsequently transmitted via consumption of contaminated fresh water.

It has been proposed that cholera outbreaks and *V. cholerae* populations are inversely correlated with the phage levels in the aquatic environment (11, 12), and a predation role was suggested for JSF4 (9), an ICP1-like phage (25). Our results do not support this model, since ICP2 and ICP3 were shown to be incapable of preying on *V. cholerae* in fresh or estuary water, and ICP1 was incapable of preying on *V. cholerae* in fresh water. However, since ICP1 is capable of predation in estuary water, it might play a role in reducing populations of *V. cholerae* is estuarine environments.

We also found that ICP1 is able to prey on stationary phase or non-dividing *V. cholerae*, perhaps suggesting a role for predation in the long-term maintenance of ICP1 in nutrient-poor aquatic reservoirs. This may be a factor in the high frequency of isolation of ICP1 in patient stools and aquatic environments in Bangladesh (9, 15). ICP1 shares features with *Escherichia coli* T4 phage (15) and it was shown that T4 has the ability to kill stationary phase *E. coli* (26). However, we do not yet know how ICP1 infects stationary phase *V. cholerae*. It is known that genome injection by T4 requires a membrane potential (27). However, *V. cholerae* was highly motile, which requires a membrane potential (31), even after 48h in fresh water in the absence of added carbon source. Thus, absence of a membrane potential is unlikely to explain the lack of killing by ICP1 in this setting. Our results indicate that the presence of NaCl could be an important factor for ICP1 infection. This could be due to the Na^+^ which has been shown to be important for lambda DNA injection (28).

It remains possible that ICP1 can prey on *V. cholerae* in fresh water due to the complex Bangladeshi climate scenario with high temperatures in summer that evaporate fresh water increasing its salinity in addition to the monsoon and flooding that leads to the mixture of aquatic reservoirs. Another possible scenario is that ICP1 in fresh water could be ingested in planktonic form or when attached to *V. cholerae* and also replicate within the host as has been suggested (9).

ICP2 has been found sporadically in patient and environmental samples (15, 29). Our results show that ICP2 was mostly inefficient in attaching to *V. cholerae* and had modest predation capability in the presence of chitin in estuary conditions. ICP2 seems to thrive instead within the host and nutrient-rich environments.

The last phage we studied was ICP3, a T7-like phage (15). We show that ICP3 is able to attach to *V. cholerae* but is incapable of replicating in fresh or estuary water, and the majority becomes inactive after a short period of time. ICP3 when bound to outer membrane vesicles cannot inject its DNA and 90% of bound ICP3 shows a full capsid (30). T7 DNA ejection needs active transport into the bacterial host that is mediated by the host RNA polymerase (31). *V. cholerae* remained highly motile in the nutrient poor settings we tested, suggesting it is metabolically active, but it is possible that the host RNA polymerase is not available to aid in phage DNA transport into the cell. In spite of the majority of ICP3 becoming inactive after adsorption to *V. cholerae*, a fraction of the phage remained intact as phage were recovered in the majority of infected animals. This could explain why it is found sporadically in cholera patient samples and suggests that ICP3, like ICP2, has adapted to replicate within a nutrient-rich microenvironment.

Because of ICP3’s fickle nature in water, it is possible that maintenance of ICP3 in Bangladesh relies primarily upon rapid transmission between people such as occurs within households (32). Of 29 sequences reported for environmental *V. cholerae* phages (11, 25, 29, 33), 41% corresponded to ICP3-like phages. This result is striking, since our model does not support the notion that ICP3 can replicate in nutrient-limiting aquatic environments. However, detailed information about the exact locations for sampling is lacking in these studies. It is possible that these phages were collected from surface waters near fish markets or nutrient-rich microenvironments that allow *V. cholerae* growth (34) or near sewage where human-shed phage could be collected.

ICP2 and ICP3 also fared poorly at predation in the infant mouse small intestine after being co-inoculated in cell-associated form. This contrasts with the sporadic occurrence of these phages, often at high titers, in cholera patient stools (15, 25). The inability of these phage to replicate in our animal experiments could be due to any number of reasons. For example, niches that support replication within the human small intestine may not be reproduced in infant mice. Alternatively, replication of *V. cholerae* in the absence of associated phage is required to occur for some period of time before free phage can successfully initiate replication. *V. cholerae* escape mutants have been observed in patient stools with high titers of ICP2 (35). The absence of ICP2 escape mutants in our mouse colonization experiment is consistent with a lack of sufficient phage predation needed to impose selection on *V. cholerae*.

Our results suggest that ICP2 and ICP3 evolved to replicate within nutrient rich environments, which might occur in certain environmental settings and within the human intestinal tract. On the other hand, our data suggest that ICP1 has evolved to be adaptable for predation on *V. cholerae* both in the human host and in aquatic environments given adequate salinity. These different niche adaptations may help to maintain all three phages in the complex cholera endemic setting of Bangladesh.

In summary, our work highlights the evolution of niche-specific predation by three virulent phages. We examined adaptation for predation in different environments including the small intestine and aquatic environments that vary by Na^+^, carbon and nitrogen source concentrations. Our results suggest that virulent phages have evolved to infect *V. cholerae* at different stages of this pathogen’s life cycle to ensure their survival. This study will improve our understanding of how cholera outbreaks are impacted by the negative selection pressures of virulent phages. This will ultimately help researchers to develop more accurate models for predicting the dynamics of cholera outbreaks, as well as enlighten the possible use of virulent phages to combat this disease.

## Materials and Methods

### Strains, media and growth conditions

*V. cholerae* HC1037, a O1 El Tor clinical isolate (30) was grown in LB Miller broth or LB Miller agar plates at 37°C. Fresh water was collected from Massapoag Lake in MA, USA. Its chemical composition is similar to pond water obtained from a cholera endemic area in Bangladesh (10, 17, 18, 36, 37). Estuary conditions were emulated by a 7 g/L (0.7%) solution of Instant Ocean^®^ Sea Salts (Spectrum Brands, Inc). Phages used in this study were isolated from patient samples from Bangladesh and correspond to: ICP1 (ICP1_2011_A; Myoviridae), ICP2 (ICP2_2004; Podoviridae) and ICP3 (ICP3_2007; Podoviridae) (15).

### Phage adsorption and predation assay in aquatic microcosms

Stationary phase *V. cholerae* were diluted 1:1000 and grown for 8h at 37°C with aeration. Then, 5 µl (∼10^8^ CFU) were inoculated into 2.5 ml of autoclaved, filter sterilized fresh water or 0.7% instant ocean and incubated overnight at 30°C with aeration for adaptation. In some experiments, 1% shrimp chitin flakes (Sigma-Aldrich) was added and cultures were pre-adapted statically at 30°C.

After adaptation, phage were added at an MOI of 0.01. Adsorption was measured after static incubation at 24°C for 15 min. Phage predation was assessed at 30°C with aeration: After 1, 2, 4, 7 and 24h, aliquots were taken to measure bacterial viability by plating and free phage in the supernatant by plaque assay as described in the supplemental information. Survival rate of *V. cholerae* was calculated as follows: CFU/ml infected /CFU/ml initial and converted logarithmically.

To measure phage replication, infected cultures were centrifuged to separate bacteria from free phage in the supernatant. 500 µl of supernatant were filter sterilized using a 0.22 μm filter (COSTAR). The titer of phage was measured by plaque assay.

To measure viable phage associated with *V. cholerae*, infected cells were washed in either fresh water or 0.7% Instant Ocean. Bacterial suspensions were serially diluted, fresh *V. cholerae* was added and mixtures were allowed to incubate for 10 min at 24°C. Each dilution was transferred to 24-well clear, for plaque assays.

Phage stability was measured by plaque assays after incubating each phage for 24h in either fresh water or 0.7% Instant Ocean in the absence of a host strain. All samples were analyzed using one-sample t-test and GraphPad Prism version 6.00.

### Infant mice colonization

To evaluate the impact of ICP2 or ICP3 on colonization, *V. cholerae* pre-adapted overnight in fresh water were incubated with each phage at an MOI of 1 or in fresh water alone. After 45 min of incubation, bacteria were washed to remove unbound phage and 50 µl containing ∼10^5^ CFU were inoculated orogastrically to 5-day-old litters (both sexes) of CD1 mice (38). 24h post infection, mice were euthanized and small intestines were collected for homogenization in 1 ml of LB broth supplemented with 20% glycerol. Bacterial load was assessed by serial dilution and plating of viable CFU on LB agar plates supplemented with Streptomycin 100 µg/ml (36, 39). PFU were enumerated by plaque assay. Results were analyzed using one-sample t-test and GraphPad Prism version 6.00.

Animal procedures were conducted in accordance with the rules of the Department of Laboratory Animal Medicine at Tufts University School of Medicine.

## Supporting information

## Acknowledgments

We thank D. Lazinski and R. Molina-Quiroz for reviewing this manuscript and the members of the lab for helpful discussions. This work was supported by the Howard Hughes Medical Institute and NIH grant AI055058 to A.C. C.A.S.V. was supported by the Pew Latin American Fellows Program in the Biomedical Sciences from PEW Charitable trusts and a CONICYT Becas Chile postdoctoral fellowship.

