## Supplementary material for "Niche adaptation limits bacteriophage predation of *Vibrio cholerae* in a nutrient-poor aquatic environment"

**This PDF file includes**

Supplementary Materials and Methods
Figs. S1 to S6
Table S1
References for SI reference citations

Cecilia A. Silva-Valenzuela, Andrew Camilli

### SI Materials and Methods

#### Bacteriophage stock preparation

For high-titer phage stocks, 40 ml cultures of *V. cholerae* HC1037 were grown to mid-exponential phase ($OD_{600}$:0.1-0.2) and inoculated with 5 fresh plaques from each phage. Infected cultures were incubated at 37°C with aeration for 1.5 hours. After this time, sodium citrate was added at a final concentration of 50 mM for 1 h to reduce further phage adsorption. After visualization of lysis, cultures were centrifuged to spin down debris and intact cells and the supernatant was filter-sterilized using a 0.22 µm filter (Millipore). A 0.2 volume of 5x phage precipitation buffer (20% Polyethylene glycol MW 8000, 2.5 M NaCl) was added and mixed by inversion. The solution was incubated at -80°C for 20 minutes and thawed at 4°C to precipitate phage. Finally, phage were concentrated by centrifugation at 4°C (10,000 RCF, 10 min), supernatant was removed and the pellet was resuspended in STM buffer (100 mM NaCl, 10 mM MgSO4, 10 mM Tris-HCl pH 7.5). Phage stocks were stored at 4°C until further use.

#### Titering phage concentration by plaque assay

Ten microliters of serially diluted phage was mixed with 100 µl of OD=0.1 *V. cholerae* HC1037 and phages were allowed to adsorb for 10 min at 24ºC. Each dilution was transferred to 24-well clear, untreated tissue culture plates (Corning) and 500 µl of molten 50˚C 0.3% agarose in LB Miller broth was added and mixed by gentle swirling. Plates were incubated overnight at 37˚C and plaques were counted.

Cecilia A. Silva-Valenzuela, Andrew Camilli

**Analysis of phage escape mutants**

After phage predation assays in either fresh water or 0.7% instant ocean, single *V. cholerae* cells that survived predation were colony purified on LB plates. Individual colonies were exposed to the corresponding phage by cross-streaking (1). Briefly, a 20 µl aliquot of phage at $10^9$ PFU/ml was spotted onto the edge of an LB plate, and the plate was tipped at an angle to allow the dribble across the center line of the plate. After letting the liquid soak into the plate, individual colonies of *V. cholerae* were streaked across the line of phage, and the plate was incubated overnight at 37˚C. Sensitivity to the phage was revealed by no or poor growth after going through the line of phage.

Resistant and sensitive mutants were grown overnight at 37˚C with aeration. Genomic DNA was extracted using the DNeasy Blood & Tissue Kit (Qiagen). The extracted DNA was used to prepare whole-genome libraries using the Nextera XT DNA Library Preparation Kit (Illumina). Samples were sequenced as single-end 50-bp length on an Illumina HiSeq 2500. Resulting reads were compared to the reference genome for *V. cholerae* HC1037 using the CLC Genomics Workbench 8 software (Qiagen) and variant analyses were performed on mapped reads with a frequency threshold of 51%.

Cecilia A. Silva-Valenzuela, Andrew Camilli

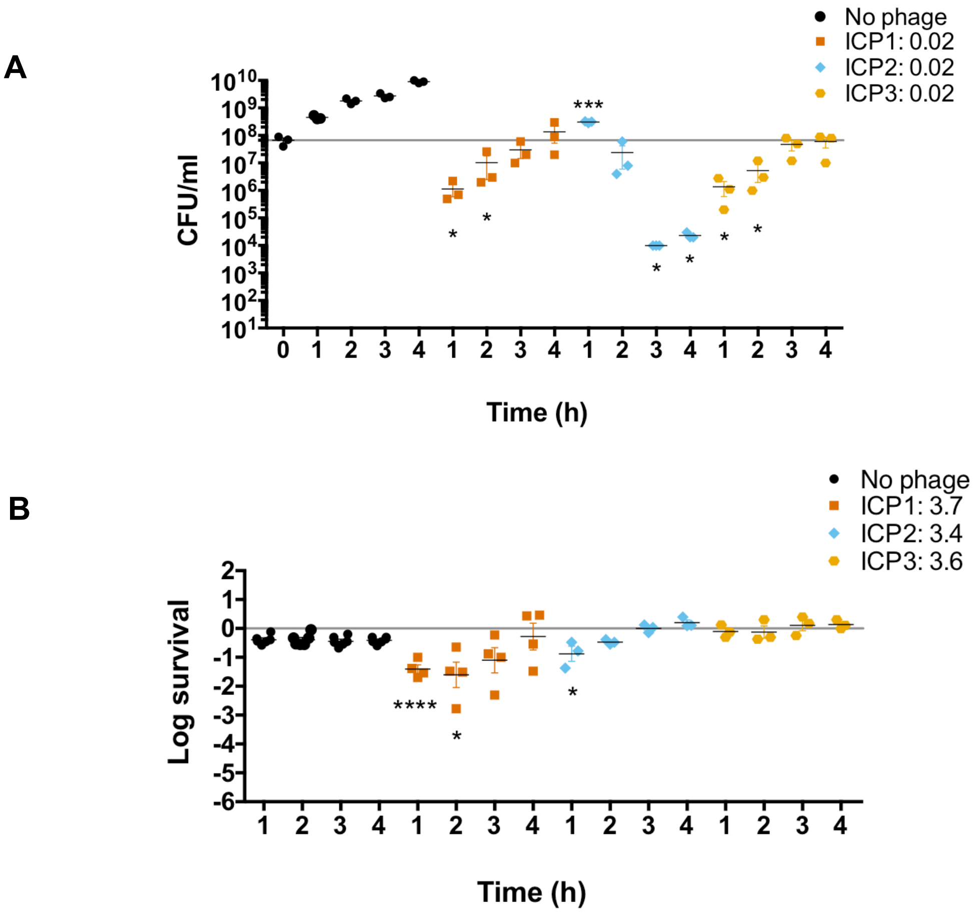


**Figure S1. ICP phage predation dynamics on rapidly growing and stationary phase bacteria.** (a) $10^6$ CFU of *V. cholerae* from an overnight culture were inoculated into M9 minimal media in the absence of added carbon source. Then each ICP phage was added to an MOI> 3 to assess bacterial viability over time. Graph represents the average and standard deviation of at least three biological replicates. (b) Cultures were grown in LB broth at 30ºC with aeration until they reached $OD_{600}$ of 0.1. Then, phages were added to an MOI of 0.01 to assess bacterial viability over time. Graph represents the average and standard error of at least three biological replicates. Grey line indicated bacterial concentration at time 0. (*P<0.05, ***P<0.001, ****P<0.0001).

Cecilia A. Silva-Valenzuela, Andrew Camilli

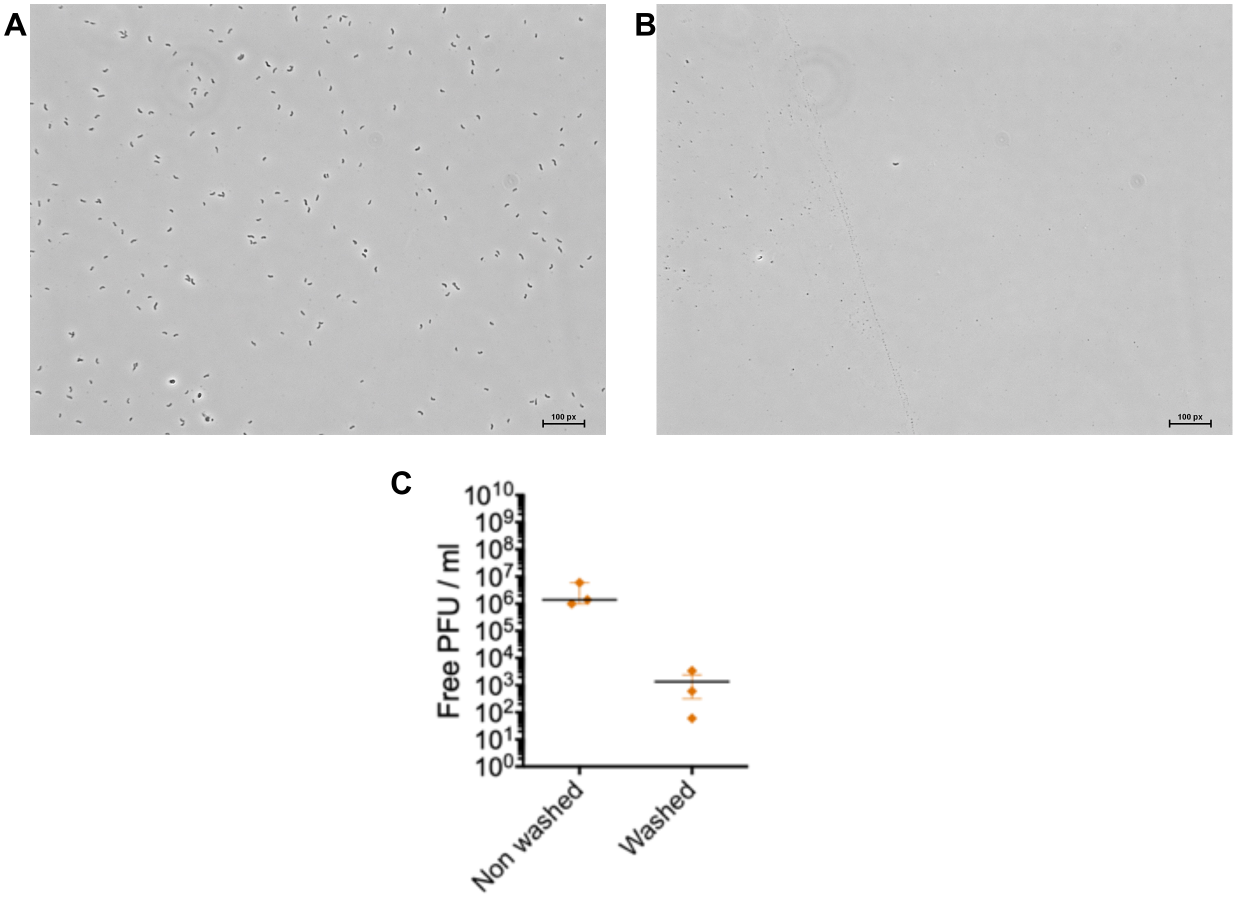


**Figure S2. ICP1 preys on V. cholerae in estuary water**. 107 CFU of V. cholerae from an overnight culture were pre-adapted in 0.7% Instant Ocean overnight in the absence of added carbon source. ICP1, ICP2 or ICP3 were added to an MOI of 0.01 to assess bacterial viability and phage replication over time. Phase contrast microscopic images of non-infected control cultures (a) and infected cultures (b) after 6h of ICP1 infection. Scale bar represents 65 µm (0.65 µm/px) (c) Free PFU titer after 6h of infection (Non-washed). After washing the infected cells, samples were filtered using a 0.22 um filter to test for free PFU. Graph shows median with range of three biological replicates.

Cecilia A. Silva-Valenzuela, Andrew Camilli

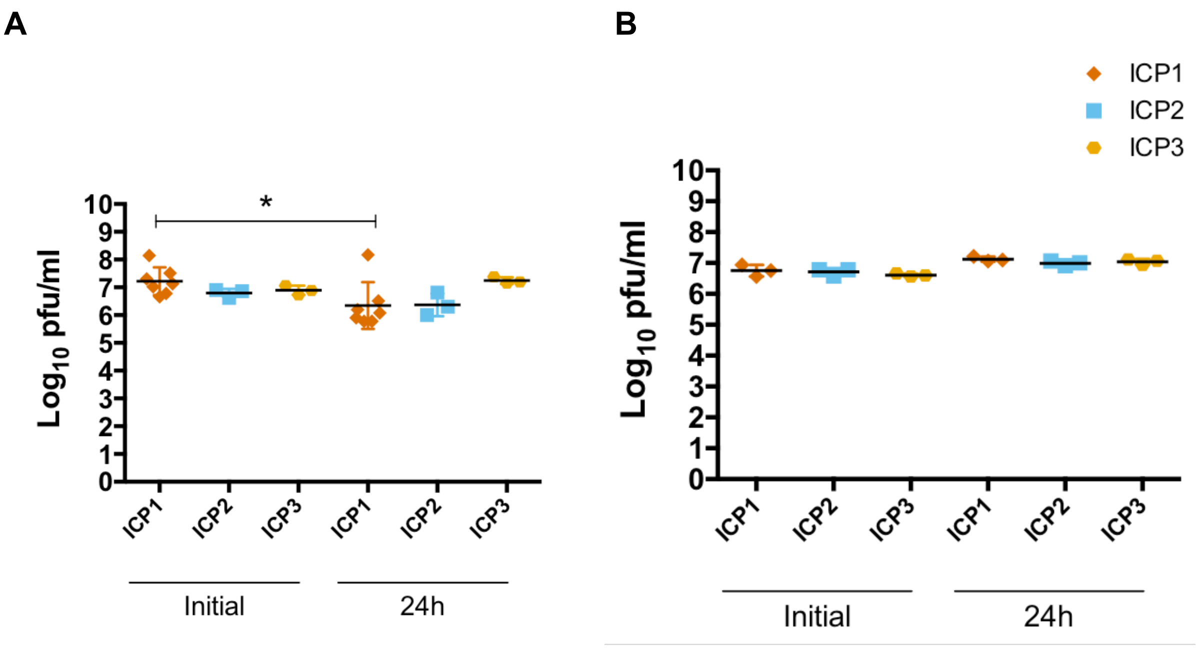


**Figure S3. Stability of ICP phages in estuary or fresh water.** $10^7$ PFU of ICP1, ICP2 or ICP3 were added to (a) estuary or (b) fresh water in the absence of a bacterial host. PFU/ml were measured at 30°C with aeration after 24h.

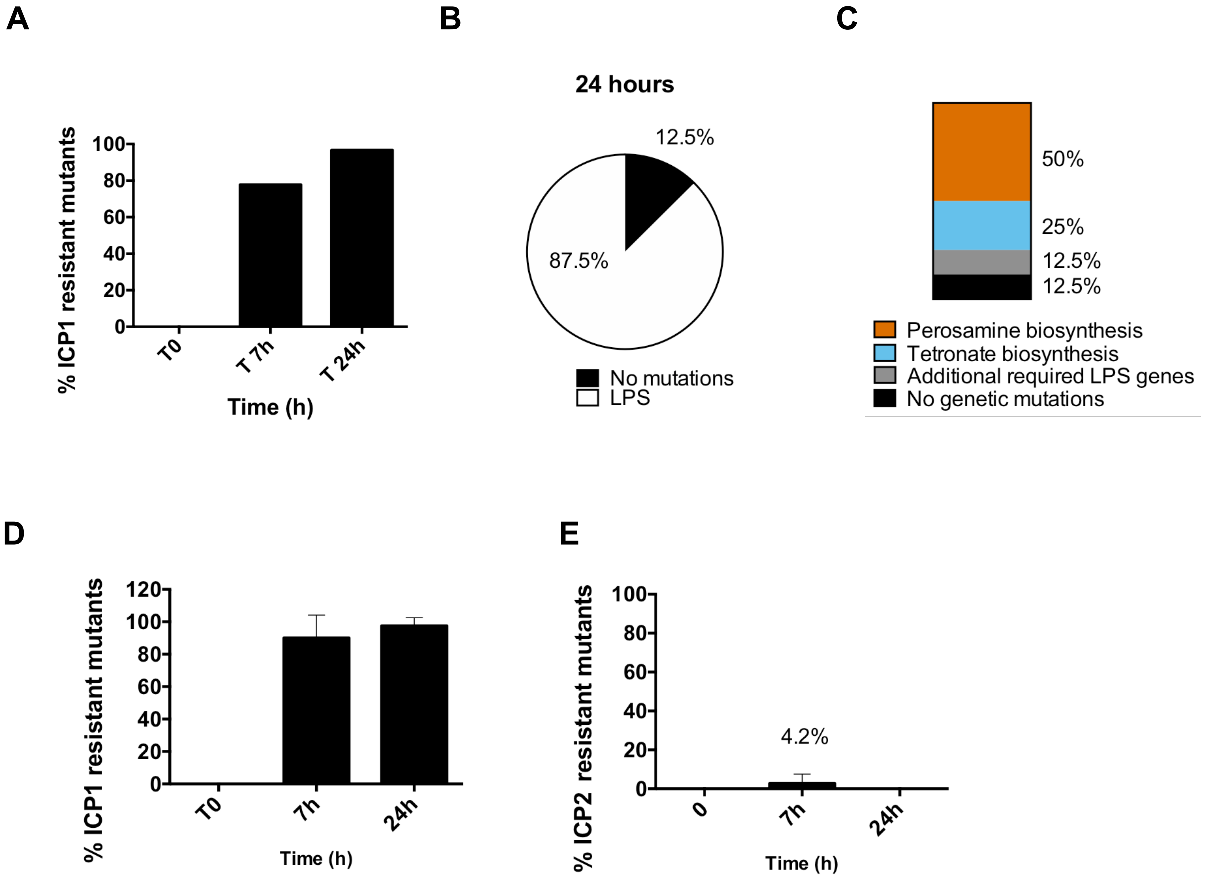


**Figure S4. Phage escape mutants carry genetic mutations linked to the phage receptor**. Percentage of survivors resistant to ICP1 or ICP2 after predation in estuary conditions or in the presence of 1% chitin. (a) Number of ICP1 resistant mutants over time. (b) Genetic mutation linkage. (c) Detailed information of pathways where genetic variations were found on ICP1 resistant mutants. (d) Number of ICP1 resistant mutants over time in 1% chitin in estuary conditions. (d) Number of ICP2 resistant mutants over time in 1% chitin in estuary conditions.

Cecilia A. Silva-Valenzuela, Andrew Camilli

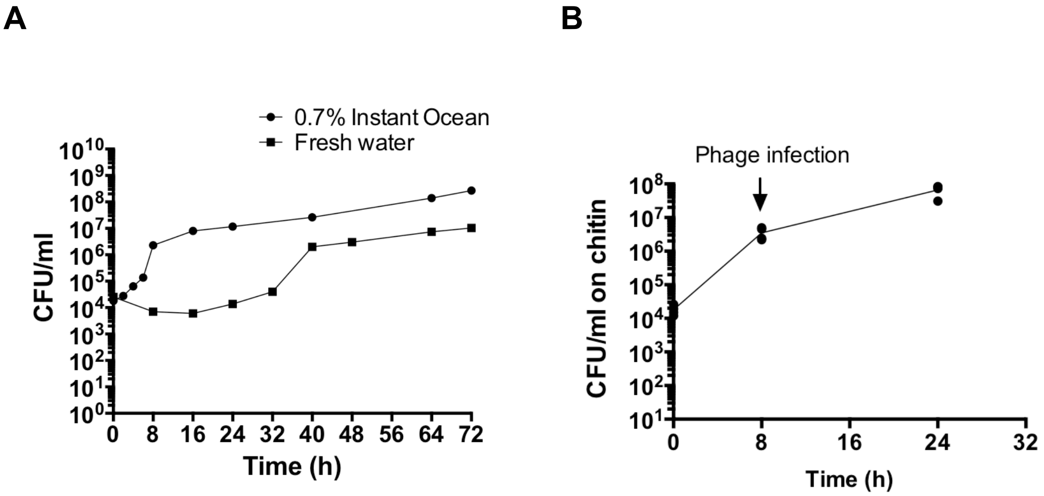


**Figure S5. *V. cholerae* grows in the presence of chitin in fresh water and estuary environments**. $10^4$ CFU of *V. cholerae* from an overnight culture were inoculated into a (a) 1% chitin suspension in 0.7% Instant Ocean or autoclaved, filter-sterilized fresh water. Cultures were incubated statically at 30°C and vigorously shaken before every time point. Graph represents the average and standard error of at least three biological replicates. (b) Bacterial numbers in 1% chitin on 0.7% Instant Ocean at 0, 8 and 24h in 1% chitin for phage predation assay.

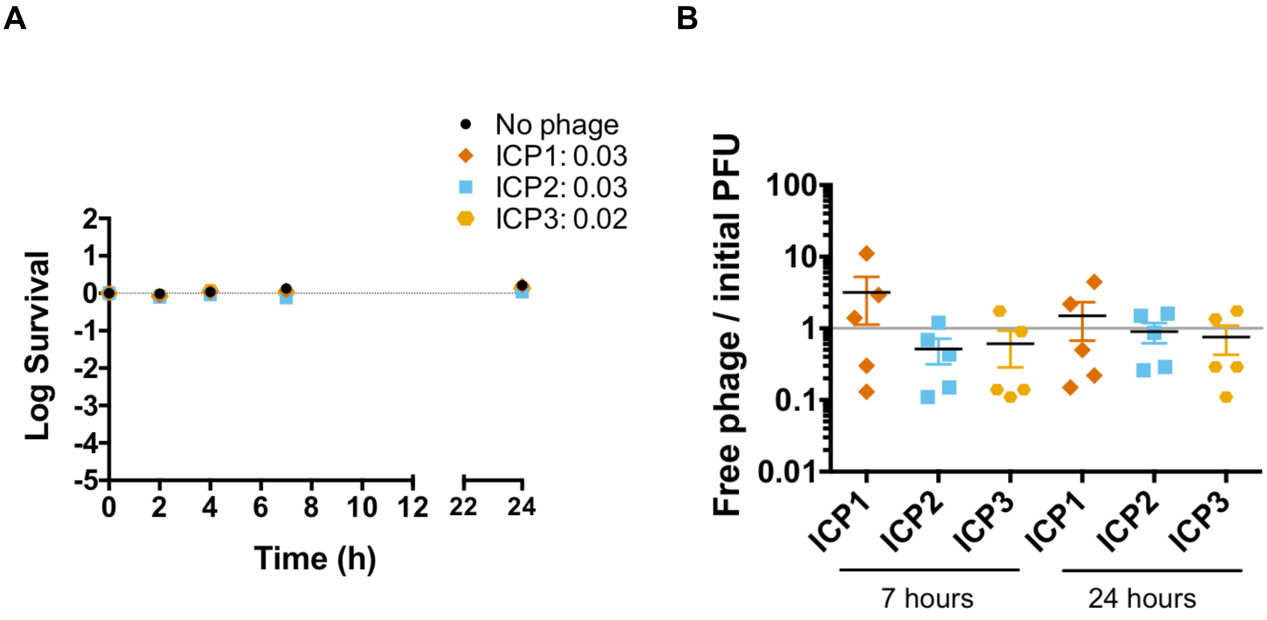


**Figure S6. Chitin availability does not aid phage predation in a nutrient-poor fresh water environment.** $10^7$ CFU of *V. cholerae* from an overnight culture were pre-adapted in a 1% chitin suspension in autoclaved, filter-sterilized fresh water. The next day, phages ICP1, ICP2 or ICP3 were added to an MOI of 0.01 to assess bacterial viability and phage replication over time. Graph represents the average and standard error of at least four biological replicates. (a) Bacterial viability at 1, 2, 4, 7 and 24h in fresh water. (b) Phage replication at 7 and 24h in fresh water relative to number initially added. Grey line indicates initial phage added.

**Table S1: Genetic variations in ICP1 resistant survivors in planktonic 0.7% Instant Ocean at 24h post-infection**

| Isolate | Reference Position | Gene | Annotation | Type | Reference | Coverage | Frequency | Amino acid change |
|---|---|---|---|---|---|---|---|---|
| ICP1$^{R}$ 1 | 2663871 | *wbeL* (A track) | | Deletion | A | 41 | 82.93 | WP_000117645.1:p.Thr39fs |
| ICP1$^{R}$ 2 | 2658706 | *manA* (A track) | GDP-mannose 4,6-dehydratase | Deletion | A | 28 | 82.14 | WP_001036868.1:p.Thr127fs |
| ICP1$^{R}$ 3 | 2672896 | *wbeV* | Glycosyl transferase family 1 (RS112245) | SNV | A | 42 | 100 | WP_000865953.1:p.Leu74Arg |
| ICP1$^{R}$ 4 | 2660080 | *wbeE* | aminotransferase DegT (RS12185) - Perosamine synthase | Insertion | - | 26 | 65.38 | WP_000613529.1:p.Asn207fs |
| ICP1$^{R}$ 5 | 2657695 | *manB* | phosphomannomutase (RS12175) | SNV | T | 50 | 90 | WP_000661577.1:p.Asp252Glu |
| ICP1$^{R}$ 6 | 2656372 | *manC* | mannose-1-phosphate guanyltransferase (RS12170) | SNV | G | 74 | 100 | WP_001894734.1:p.Trp278Leu |
| ICP1$^{R}$ 7 | No genetic variations | | | | | | | |
| ICP1$^{R}$ 8 | 2664290 | *wbeL* (no A-track) | | SNV | C | 17 | 100 | WP_000117645.1:p.Pro176Leu |
